# Allosteric modulation of GABA_A_ receptors in rat basolateral amygdala blocks stress-enhanced reacquisition of nicotine self-administration

**DOI:** 10.1101/2020.07.10.197525

**Authors:** Burt M Sharp, Qin Jiang, Xenia Simone, Petra Scholze

**Affiliations:** Department of Genetics, Genomics and Informatics, College of Medicine, University of Tennessee Health Science Center, Memphis, Tennessee 38163; Department of Pathobiology of the Nervous System, Center for Brain Research, Medical University of Vienna, Vienna, Austria

**Keywords:** allosteric, GABA_A_, basolateral amygdala, stress, nicotine, relapse

## Abstract

Stress is a major determinant of relapse to smoked tobacco. In a rat model, repeated stress during abstinence from nicotine self-administration (SA) results in enhanced reacquisition of nicotine SA, which is dependent on the basolateral amygdala (BLA). We postulate that repeated stress during abstinence causes hyperexcitability of BLA principal output neurons (PN) due to disinhibition of PN from reduced inhibitory regulation by local GABAergic interneurons. To determine if enhanced GABAergic regulation of BLA PNs can lessen the effects of stress on nicotine intake, positive allosteric modulators (PAMs) of GABA_A_ receptors were infused into the BLA immediately prior to reacquisition of nicotine SA. Three selective PAMs (e.g., NS 16085, DCUK-OEt, DS2) with varied GABA_A_ subunit specificities abolished the stress-induced amplification of nicotine taking during reacquisition. These studies indicate that highly selective PAMS targeting α3 or δ subunit-containing GABA_A_ in BLA may be effective in ameliorating the stress-induced relapse to smoked tobacco during abstinence from cigarettes.

## Introduction

Nicotine is the principal psychoactive agent in smoked tobacco. My laboratory developed the operant behavioral paradigm for virtually unlimited (i.e., 23 hour) access to nicotine self-administration (SA) in order to model human smoking (1). More recently, we modeled the effects of stress during abstinence from nicotine SA, which is among the most prevalent causes of relapse to smoking. We have shown that repeated stress during abstinence from nicotine SA results in enhanced nicotine SA and enhanced motivation to obtain nicotine following abstinence (2). As such, this is a model for stress-induced relapse to smoked tobacco in humans. Additional experiments showed that circuitry in the basolateral amygdala (BLA) and its connections to nucleus accumbens core are necessary for the expression of stress-enhanced nicotine SA (3).

We postulate that repeated stress during abstinence causes hyperexcitability of BLA principal output neurons (PN) due to disinhibition of PN from reduced inhibitory regulation by local GABAergic interneurons (4, 5). These interneurons activate GABA_A_ receptors on BLA principal neurons (PN) that project to multiple brain regions, including monosynaptic glutamatergic efferents to median spiny neurons in the nucleus accumbens. GABA_A_ receptors are ligand-gated pentameric chloride channels principally assembled from α, β, and γ or δ subunits (6). BLA PNs express a variety of GABA_A_ receptors (GABA_A_) containing α2, α3, β, γ2 and δ subunits (7–9).

Based on the foregoing hypothesis, we determined if enhanced GABAergic regulation of BLA PNs can lessen the effects of stress on nicotine intake. Positive allosteric modulators (PAMs) of GABA_A_ were infused into the BLA immediately prior to the reacquisition of nicotine SA by rats that had previously been stressed during abstinence from nicotine. PAMs targeting different GABA_A_ subtypes present in BLA were evaluated: NS16085 (4-chloro-3-{6-[5-(2-hydroxypropan-2-yl)-1H-1,3-benzodiazol-1-yl]pyridin-2-yl}benzonitrile) is highly specific for the benzodiazepine binding site on α2/α3 subunit-containing GABA_A_ (10); DCUK-OEt (5,7-dichloro-4-([diphenyl carbamoyl] amino) quinoline-2-ethyl carboxylate) binds to a novel, benzodiazepine and flumazenil insensitive GABA_A_ site that requires specific subunit combinations (i.e., α1 or α5, or β2 or β3, γ2 or δ) (11); and DS2 (4-chloro-N-[2-(2-thienyl)imidazo[1,2-a]pyridine-3-yl benzamide) binds exclusively to δ subunit-containing GABA_A_ (12, 13).

We found that all three GABA_A_ PAMs diminished the stress-induced amplification of nicotine intake following abstinence from the drug. This suggests that more than one subtype of GABA_A_ modulates BLA neuronal activity, which is affected by chronic stress during abstinence from nicotine. Highly selective PAMs, such as DCUK-OEt, may be beneficial in reducing stress-related relapse to cigarette smoking.

## Results

To verify the location of microinfusions into BLA, random animals that had previously been implanted with guide cannulae were injected with AAV-CAMKII-mCherry; brains were obtained after completion of behavioral studies. Figure 1 demonstrates the diffusion and expression throughout BLA of a representative infusion of AAV-CAMKII-mCherry. Fluorescence is largely confined to and detectable throughout most of BLA. Figure 2 shows BLA schematics of injection sites in rats infused with each PAM.

**Figure 1.**
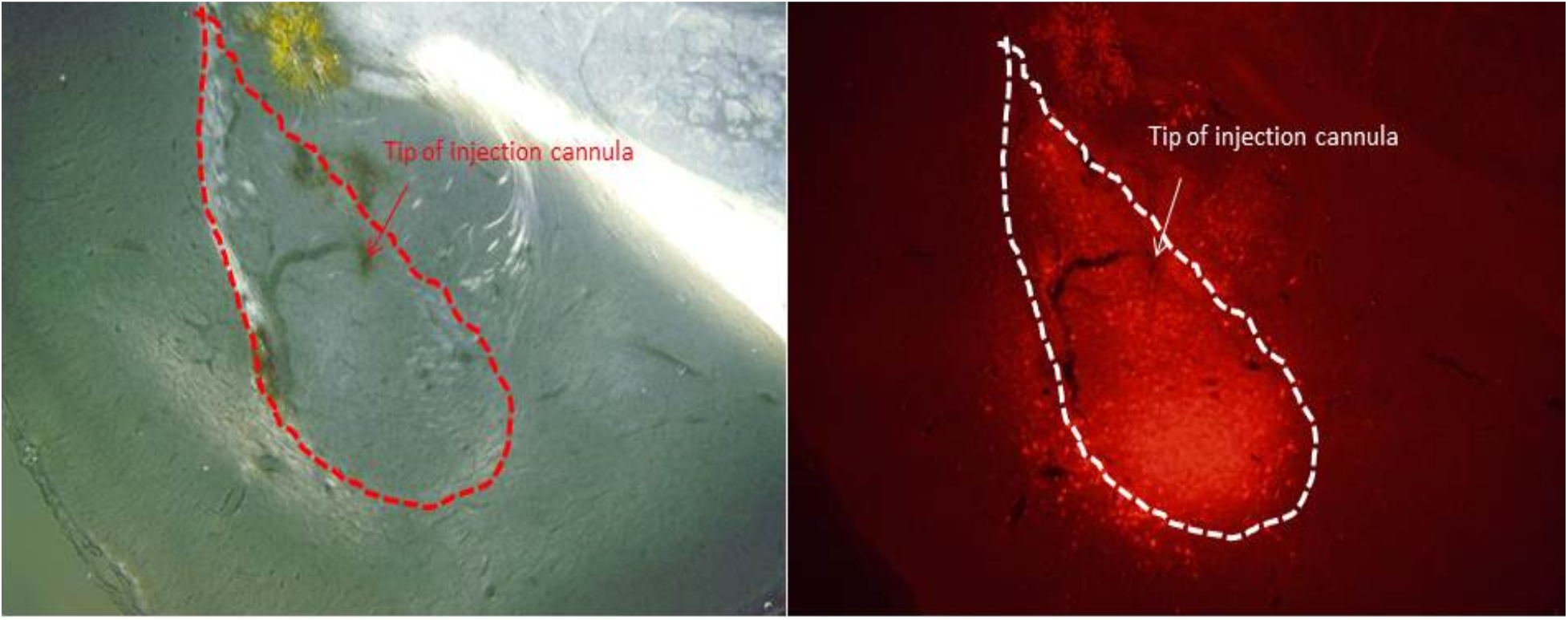
Photomicrographs of AAV-CAMKII-mCherry expression in BLA: (L) BLA (DIC; AP −2.8, ML ±4.6, DV −7.4) shows small amount of old hemorrhage (yellow) surrounding tip of implanted guide cannula just above BLA; tip of injection cannula is shown by arrow; (R) mCherry fluorescence expressed in BLA neurons.

**Figure 2.**
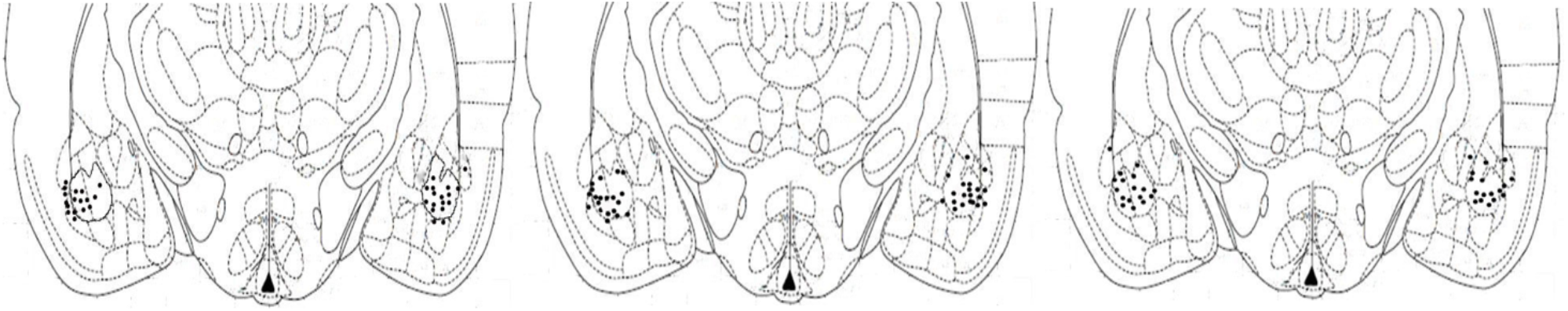
BLA schematics of injection sites in rats infused with NS 16085 (L), DCUK-OEt (middle) or DS2 (R).

Figure 3 shows the effects of bilateral BLA infusions of NS16085 (20 μM) on the reacquisition (ReAq) of operant nicotine SA after chronic exposure to stress during abstinence. Reacquisition of nicotine SA was assessed during three consecutive daily sessions, beginning 24h after the 4th stress. NS16085 (20 μM per side) or vehicle was infused 30 min before nicotine SA sessions on d1 and d2 of reacquisition (arrows). In the group receiving vehicle, active lever presses and nicotine infusions increased after repeated stress on day 1 of reacquisition compared to acquisition (Acq) (F=10.3, p=0.006 and F=3.5, p=0.080 for active lever and infusion, respectively). The interaction of stage [reacquisition vs acquisition] x treatment was significant for both lever and infusion (F=11.8, p=0.004 and F=10.1, p=0.006 for lever and infusion, respectively). In the vehicle group, active lever presses and infusions were both significantly greater on reacquisition day 1 compared to acquisition (*p* < 0.00006; Tukey). Additionally, on reacquisition day 1, treatment with NS 16085 blocked the enhancing effects of repeated stress on both lever presses and infusions observed in the vehicle group (p < 0.003; Tukey).

**Figure 3.**
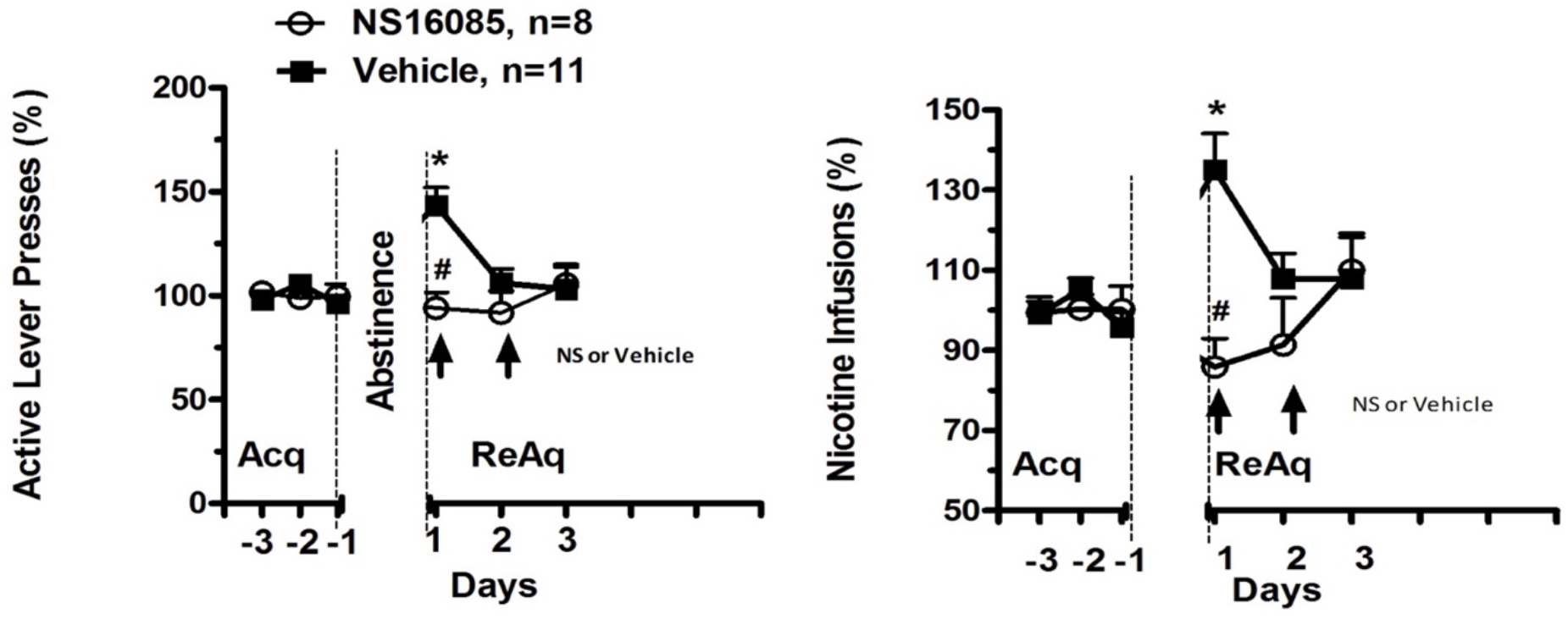
Effect of bilateral BLA infusion of NS16085 on the reacquisition (ReAq) of nicotine SA after chronic exposure to stress during abstinence. Reacquisition of nicotine SA was assessed 24h after 4th stress. NS16085 (20 μM) was infused into BLA 30m before nicotine SA sessions on d1 and d2 of ReAq. Active Lever and Nicotine Infusion: Stage [Acquisition (Acq) v ReAq] and Stage x Treatment (repeated measures ANOVA): F=10.3, p=0.006 and F=11.8, p=0.004, respectively for lever; F=3.5, p=0.080 and F=10.1, p=0.006, respectively for infusion. \**p* < 0.00006 v Acq within group, # p < 0.003 NS v vehicle at d1 (Tukey).

Figure 4 shows the effects of infusing DCUK-OEt (5 or 20 μM) or vehicle into BLA bilaterally 30 min before nicotine SA sessions on d1 and d2 of reacquisition (ReAq, arrows). Chronic stress increased active lever presses and infusions in the groups receiving vehicle or DCUK-OEt 5 μM during reacquisition, but not in the DCUK-OEt 20 μM group (2-way ANOVA: F=13.7, p=0.004 and F=14.1, p=0.002 for lever and infusion, respectively). A significant interaction of stage x treatment was also found for active presses and infusions (F=15.2, p=0.003 and F=18.3, p=0.0009 for lever and. Infusions, respectively). Lastly, on d 1 of reacquisition, DCUK-OEt (20 μM) significantly reduced both lever presses and infusions compared to vehicle (#, p < 0.001; Tukey), without affecting the underlying operant nicotine SA. Thus, enhanced activation of BLA GABA_A_ induced by the novel PAM, DCUK-OEt, blocked the stress-enhanced component of nicotine SA.

**Figure 4.**
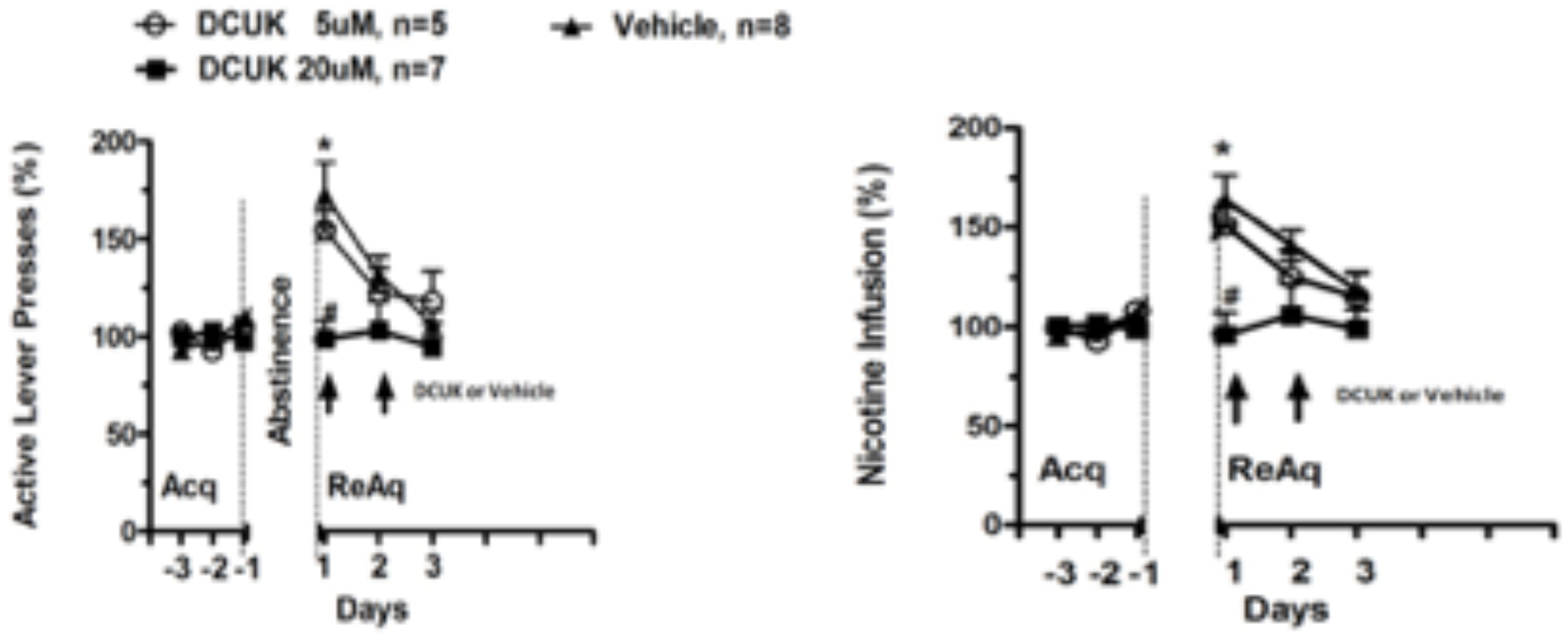
Effect of bilateral BLA infusion of DCUK-OEt on the reacquisition (ReAq) of operant nicotine SA after chronic exposure to stress during abstinence. Reacquisition of nicotine SA (FR5) was assessed 24h after 4th stress. DCUK (5 and 20 μM; 300nl/side in 5% DMSO) was infused into BLA 30m before nicotine SA sessions on d1 and d2. Active Lever and Nicotine Infusion: Stage [Acquisition (Acq) v ReAq] and Stage x Treatment (repeated measures ANOVA): F=13.7, p=0.004 and F=15.2, p=0.003, respectively for lever; F=14.1, p=0.002 and F=18.3, p=0.0009, respectively for infusion.

In Figure 5, DS2 (20 μM), a δ subunit-specific PAM, or vehicle was infused into BLA bilaterally 30 min before nicotine SA sessions on d1 and d2 of reacquisition. Chronic stress increased active lever presses and infusions in he vehicle group during reacquisition, but not after DS2 (2-way ANOVA: F=8.1, p=0.013 and F=5.7, p=0.032 for lever and infusion, respectively; *, p<0.000001 for lever and infusion in vehicle reacquisition vs acquisition, Tukey). A significant interaction of stage x treatment was also found for active presses and infusions (F=9.3, p=0.009 and F=25.5, p=0.0002 for lever and. Infusions, respectively). On days 1 and 2 of reacquisition, DS2 significantly reduced both lever presses and infusions compared to vehicle (#, p=0.009 for lever and p=0.0002 for infusion; Tukey), without affecting the underlying operant nicotine SA. Thus, enhanced activation of BLA GABA_A_ induced by the highly selective PAM, DS2, was sufficient to prevent the stress-enhanced component of nicotine SA.

**Figure 5.**
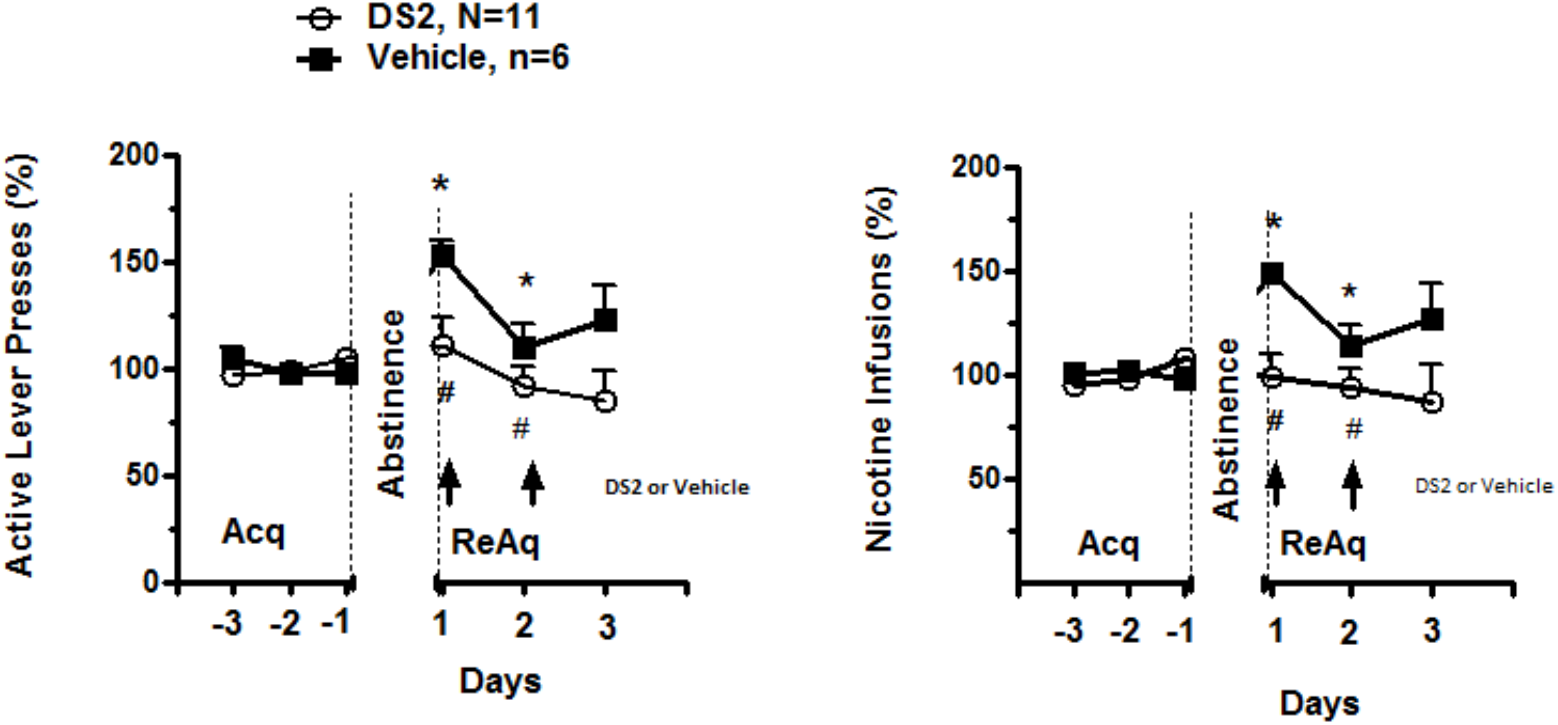
Effect of bilateral BLA infusion of DS2 on the reacquisition (ReAq) of operant nicotine SA after chronic stress during abstinence. Reacquisition of nicotine SA (FR5) was assessed 24h after 4th stress. DS2 (20 μM; 300nl/side) was infused into BLA 30m before nicotine SA sessions on d1 and d2. Active Lever and Nicotine Infusion: Stage [Acquisition (Acq) v ReAq] and Stage x Treatment (repeated measures ANOVA): F=8.1, p=0.013 and F=9.3, p=0.009 for lever; F=5.7, p=0.032 and F=25.5, p=0.0002 for infusion, respectively. *, p<0.000001 for lever and infusion in vehicle reacquisition vs acquisition (Tukey). Reacquisition Vehicle vs DS2: #, p=0.009 for lever and p=0.0002 for infusion (Tukey).

Since GABA_A_ containing α3 subunits (7) may mediate the foregoing behavioral effects of NS 16085 on BLA function, we sought to determine if these subunits are also targeted by DCUK-OEt. Electrophysiology studies have identified several α subunits that interact with this compound, although α3 was not evaluated (11). Therefore, the effects of DCUK-OEt on GABA_A_ assembled from α3 in combination with β2γ2 or β3γ2 were evaluated.

The results in Figure 6 show the potentiation by DCUK-OEt of inhibitory currents induced by GABA (EC_10_) in α3 subunit-containing recombinant rat GABA_A_ isoforms expressed by *Xenopus laevis* oocytes. As previously reported with α1β3γ2 (11), DCUK-OEt effectively and potently potentiated the inhibitory currents generated by α3β2γ2 (n=4) and α3β3γ2 (n=5): EC_50_=0.38 and 0.13 μM, respectively. This potentiation was not blocked by Flumazenil (1μM or 10μM), an antagonist specific for the GABA_A_ benzodiazepine site. Therefore, the potentiation of inhibitory currents by DCUK-OEt depends on interactions with a novel site on α3 subunit-containing GABA_A_.

**Figure 6.**
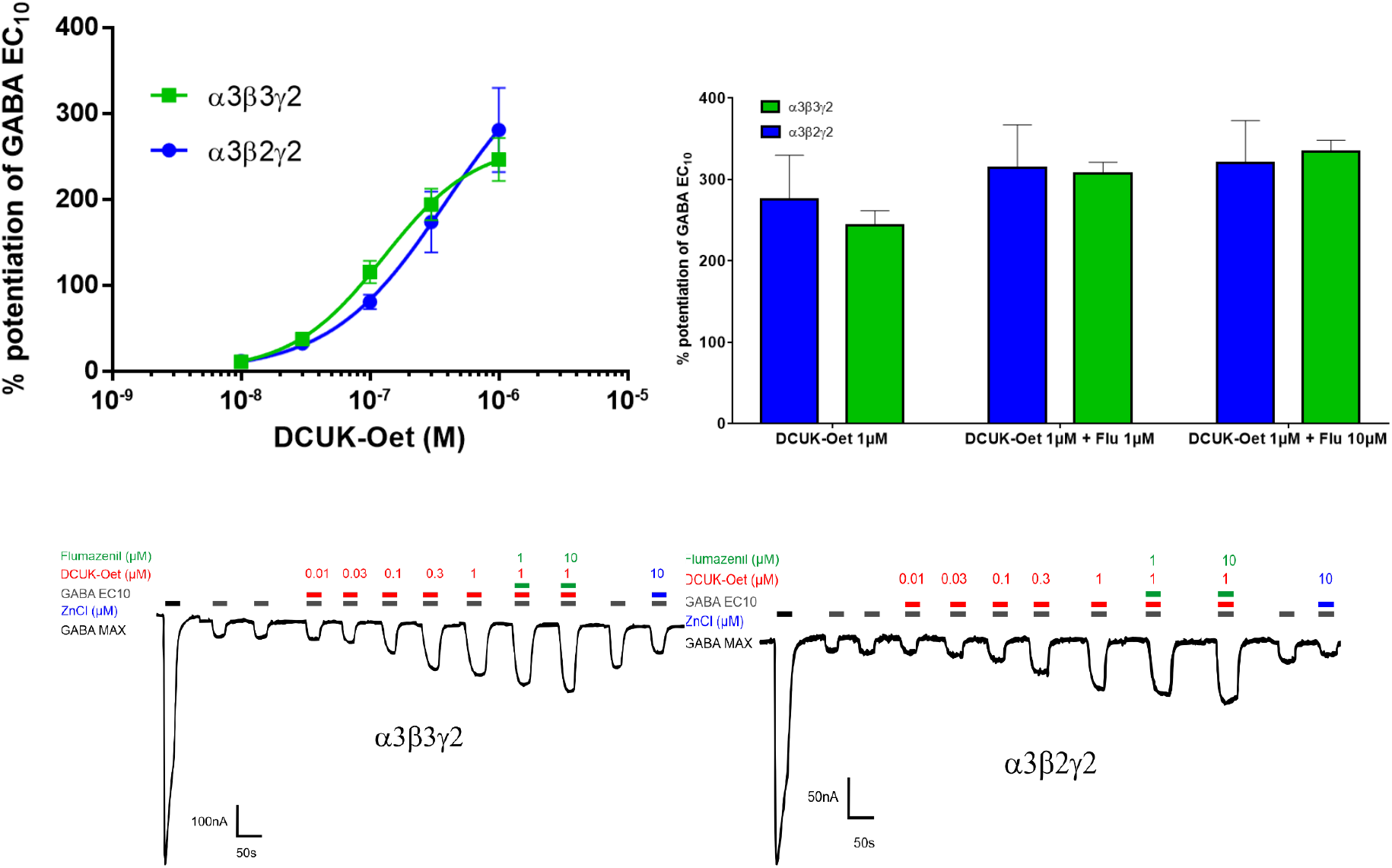
Effect of DCUK-OEt on the potentiation of α3 subunit-containing GABA_A_ inhibitory currents induced by GABA (EC_10_) in recombinant rat GABA_A_ expressed by *Xenopus laevis* oocytes. Top (L) panel shows DCUK-OEt potentiation of inhibitory currents in α3β2γ2 (n=4) and α3β3γ2 (n=5): EC_50_=0.38 and 0.13 μM, respectively. Top (R) panel shows that potentiation by DCUK-Oet 1μM was not blocked by Flumazenil 1μM or 10μM. Bottom panels show the inhibitory currents induced by GABA (EC_10_; Clamp = −70mV) and potentiated by increasing concentrations of DCUK-OEt, and the failure of Flumazenil to block this potentiation.

## Discussion

These experiments demonstrate that BLA GABA_A_ modulation by PAMs can abolish the stress-enhanced reacquisition of nicotine SA. Positive allosteric modulators, augmenting GABA_A_ activation by GABA, completely inhibited the stress-enhanced intake of nicotine and the increase in active lever presses during reacquisition of nicotine taking without altering the underlying level of either parameter achieved during the final 3d of acquisition. These observations lend support to the hypothesis that stress-enhanced reacquisition of nicotine SA involves diminished GABAergic regulation of local BLA circuits.

The following observations show that local BLA GABAergic neurons are affected by chronic stress that, in turn, alters the GABAergic regulation of BLA principal glutamatergic output neurons. These PNs, which innervate NAcc monosynaptically, are required for expression of stress-enhanced reacquisition of nicotine SA (3). 1. Dopaminergic efferents from VTA modulate a parvalbumin-positive subset of GABA interneurons within BLA (14,15), in turn, inhibit the activity of BLA PNs (7,16). 2. Extra-synaptic α3-containing GABA_A_ mediate tonic inhibition of BLA PNs; excitability of these neurons was markedly reduced by TP003, a GABA_A_ PAM that binds at the benzodiazepine site (7). 3. Stresses, such as chronic restraint, increase BLA Da release, which inhibits parvalbumin-positive GABA interneurons (17,18). Stress also sensitizes amygdaloid Da efflux to drug conditioned stimuli (19).

Taken together, the foregoing observations support the hypothesis that stress-induced increase in BLA Da decreases GABA release from parvalbumin positive BLA interneurons, thereby disinhibiting BLA PNs and promoting release of glutamate from BLA efferents to multiple brain regions. The current experimental observations are consistent with this, showing that bilateral BLA delivery of NS 16085, a highly specific PAM of α2/α3 subunit-containing GABA_A_, and DCUK-OEt, a novel PAM that interacts with several GABA_A_ including those assembled from α2 subunits (11) and α3 subunits, as shown in Fig. 6, blocked expression of the stress-enhanced component of nicotine SA during reacquisition.

BLA is regulated by two types of GABAergic currents (for review: 20). In the mouse, fast, phasic postsynaptic currents primarily occur through GABA_A_, containing α2 and some α1 subunits, but not α3 subunits. In contrast, tonic extra-synaptic inhibition, detected in BLA GABA interneurons and most principal neurons, depends largely on α3 subunit-containing GABA_A_ receptors in PNs and on non-α3 variants in interneurons (7). These tonic currents are generated by atypical non-desensitizing GABA_A_ receptors with high affinity for low levels of extra-synaptic GABA that are sensitive to benzodiazepines and TP003 (7). Blocking tonic currents increases neuronal excitability by shifting the resting membrane potential to more positive values. Conversely, amplifying the currents with TP003 produces a more negative resting potential. Thus, low concentrations of extra-synaptic GABA maintain a persistent background inhibitory conductance that affects the cellular and network E/I balance, thereby regulating intrinsic excitability and the integration of synaptic inputs (21).

There are two non-exclusive mechanisms that are likely to explain the effects of NS 16085 on stress-enhanced reacquisition of nicotine SA. NS 16085 may enhance the tonic inhibitory regulation of BLA PCs by increasing the activation of atypical extra-synaptic α3-containing GABA_A_, thereby reducing the excitability of PCs (8). Alternatively, NS 16085 may amplify the postsynaptic inhibitory currents on PCs by increasing the activity of α2-containing GABA_A_ - the primary subunit mediating these postsynaptic currents. In light of the fact that TP003, another α2/α3 subunit-specific PAM, inhibits excitability of BLA PC neurons from wild type but not α3-knockout mice, the second mechanism appears unlikely (8).

DCUK-OEt, which modulates the activity of α1-, α2-, α3- and α5-containing GABA_A_ would most likely interact with GABA_A_ that are similar to those affected by NS 16085, i.e., those containing extra-synaptic α3 subunits. In addition to modulating α3-containing GABA_A_, DCUK-OEt also might interact with α1-containing GABA_A_, although these are a minor component of the GABA_A_ controlling post-synaptic BLA PC currents; moreover, α1-containing GABA_A_ do not interact with NS 16085.

DS2 effectively blocked the stress enhanced component of nicotine SA during reacquisition. This finding implicates an additional set of GABA_A_ that are assembled from δ instead of γ subunits and are devoid of benzodiazepine binding sites. Therefore, extra-synaptic δ-containing GABA_A_ are insensitive to NS 16085. GABA_A_ that contain δ subunits mediate tonic, extra-synaptic inhibitory currents, and are usually assembled from α4 and α6 subunits (13). Modulation of these GABA_A_ by DCUK-OEt appears unlikely, since this PAM does not bind to α4δ although its interaction with α6-containing GABA_A_ is unknown (11).

In summary, three GABA_A_ PAMs were shown to block the stress-enhanced reacquisition of nicotine SA that depends on BLA (2). Based on the GABA_A_ subunit specificity of NS 16085 and DCUK-OEt, and the inhibition of BLA PN excitability by α3-containing GABA_A_, we propose that tonic, extra-synaptic α3-containing GABA_A_ are the primary BLA targets of both PAMs. The potency and efficacy of DCUK-OEt shown in our electrophysiological analyses of α3-containing GABA_A_ are consistent with this idea. The efficacy of DS2 in our behavioral model also indicates that a second subset of δ subunit-containing GABA_A_ modulate BLA excitability. These studies indicate that highly selective GABA_A_ PAMs may be effective in ameliorating the stress-induced relapse to smoked tobacco during abstinence from cigarettes.

## Materials and Methods

### Animals and surgeries

Adult male Sprague-Dawley rats (250 g, Harlan, Madison, WI) were housed in a reversed 12:12 h light/dark cycle and provided with standard rat chow and water *ad libitum*. Rats were anesthetized with ketamine/xylazine (94.3/0.11 mg/kg, respectively, i.p.) and guide cannulas (26 gauge, Plastic one, Roanoke, VA) were stereotaxically implanted bilaterally into basolateral amygdala (BLA) approximately 3 weeks prior to use. For BLA, coordinates from bregma (flat skull) are: anteroposterior (AP) = −2.56 mm, mediolateral (ML) = ±4.6, dorsoventral (DV) = −7.4 mm (22). After 2 d of recovery, jugular catheters were implanted. All procedures conformed to NIH guidelines and were approved by the Institutional Animal Care and Use Committee at the University of Tennessee Health Science Center.

### Nicotine self-administration

Rats were given extended access to nicotine SA (23 h/d, 7 d/week) without food deprivation or prior training. The nicotine SA procedure was conducted as previously described (1). Briefly, after 3 d of recovery post jugular surgery, rats were placed into an operant conditioning chamber located inside a sound-attenuating environmental chamber. The operant conditioning chamber contained two horizontal levers, and a green cue light above each lever was illuminated when nicotine was available during SA sessions. Lever presses were recorded and syringe pumps were controlled by computers and interfaces, using L2T2 software (Coulbourn Instruments, Allentown, PA, USA). Nicotine SA started immediately after initiation of the dark cycle. Pressing the active lever, randomly designated, elicited an intra-jugular injection of nicotine solution (0.03 mg/kg, in 50 μl/0.81 s, free base, pH 7.2-7.4). To avoid over-dosing from nicotine, each injection initiated a 7s timeout during which the green cue light above the lever was extinguished and nicotine was unavailable. Pressing the alternate (inactive) lever had no programmed consequence. The final hour of the 12 h lights-on cycle (i.e., 9:00–10:00 AM) was reserved for animal husbandry, refreshing the nicotine solution, and data recording. The patency of jugular catheters was checked periodically and assessed by injection of methohexital sodium (10 mg/ml, 0.1 ml; Par Pharmaceuticals, Spring Valley, N.Y.) Rats with occluded catheters were excluded from the experiment.

### Experimental design of behavioral studies

In all experiments, the protocols for acquisition of nicotine SA, abstinence, restraint stress and reacquisition of nicotine SA were conducted as previously described (2). Briefly, rats acquired nicotine SA at escalating fixed ratio (FR) schedules of reinforcement: an increasing number of active lever presses was required to receive a single injection of nicotine. Rats initiated nicotine SA on an FR 1 schedule: 1 active lever press yields 1 nicotine injection. This schedule continued for 7 d, increasing to FR 2 (for 3 d) and then FR 5 (for 7 d). After acquisition, animals were withdrawn from nicotine in their home cages (not operant conditioning chambers) for 8 d (without nicotine, levers, and cue lights). This interval was selected since the majority of attempts to quit smoking fail within the first 8 days (23) and most withdrawal symptoms, in humans and rats, peak and subside within the first week (24,25). Restraint stress was administered in a 6.5 cm diameter plastic cylinder for 30 min between 9:00–10:00 AM on d 1, 3, 5, and 7 of abstinence. 24h after the final stress, animals were returned to operant conditioning chambers for the reacquisition of nicotine SA under an FR5 schedule for a total of 3 d.

Prior to (i.e., 30 min) designated reacquisition sessions, the following compounds were infused into BLA (5 and 20μM in 0.3 μl per side over 6 min), using a microsyringe infusion pump connected to an injection cannula (33 gauge) extending 1.5 mm beyond the tip of the guide cannula: NS16085 (4-chloro-3-{6-[5-(2-hydroxypropan-2-yl)-1H-1,3-benzodiazol-1-yl]pyridin-2-yl}benzonitrile; 10), a highly α2/α3 subunit-specific positive allosteric modulator of GABA_A_ receptors; DCUK-OEt (5,7-dichloro-4-([diphenyl carbamoyl] amino) quinoline-2-ethyl carboxylate; 11), binds a novel, benzodiazepine and flumazenil insensitive GABA_A_ site requiring specific GABA_A_ subunit combinations (i.e., α1 or α5, β2 or β3, γ2 or δ); DS2 (4-chloro-N-[2-(2-thienyl)imidazo[1,2-a]pyridine-3-yl benzamide; 12,13), binds exclusively to δ subunit-containing GABA_A_; or vehicle. Injection cannulas remained in place 5 min to ensure diffusion of drug.

### Electrophysiological studies of recombinant rat GABA_A_

Recombinant rat GABA_A_ α3β2γ2 and α3β3γ2 were expressed in *Xenopus laevis* oocytes, as previously described (26). Electrophysiological studies were performed according to the methods of Borghese et al. (11). Unless otherwise stated, the washout period between the application of compounds was 3 minutes. Compounds were applied sequentially: GABA (1mM) followed by 15 min washout; GABA (EC_10_) tested twice (clamp= −70mV); DCUK-OEt at increasing concentrations (0.01-1.0 μM) + GABA (EC_10_); Flumazenil (1.0 or 10μM) + GABA (EC_10_) + DCUK-OEt (1 μM) followed by 15 min washout; Zn test to validate the incorporation of γ subunits into GABA_A_. Multiple compounds were all co-applied.

### Histology

After the final d of reacquisition, rats were anesthetized with ketamine/xylazine (94.3/0.11 mg/kg, respectively, i.p.) perfused with transcardiac paraformaldehyde, and then brains were immediately fixed in paraformaldehyde (4% in 0.1 M phosphate buffer) for 24h, dehydrated in 20% sucrose followed by 30% sucrose for 2 × 24 h, sectioned at 40μm thickness (CM1850, Leica, Nussloch, Germany), and mounted on pre-cleaned slides. The location of each microinjection cannula was determined by light microscopy.

### Data analysis and statistics

Both active lever presses and nicotine injections, during the final 3 d of acquisition of nicotine SA and during the 2 d of reacquisition of nicotine SA (FR5 schedule), were expressed as a percentage of the baseline, defined as the average number of presses or injections during the final 3 d of acquisition. The behavioral data obtained during reacquisition were analyzed by repeated measures 2-way ANOVA, using SPSS Statistics 22 (IBM, Armonk, New York), with day of reacquisition designated as a within-subjects factor and drug treatment designated as a between-subjects factor. One-way ANOVA and multiple *posthoc* comparisons (Tukey) were used to compare various time intervals within the same treatment group. Data were expressed as mean ± SEM. Statistical significance was assigned at *p* < 0.05.

## Acknowledgements

We greatly appreciate the gift of NS16085 from Karin S Nielsen at Saniona AB, Denmark. We are grateful to Lohocla Research, Inc., Aurora, Colorado for providing DCUK-OEt and partial funding to support the experiments involving this compound. We also appreciate the laboratory assistance from Guoliang Yu Ph.D.

